# NanoJ: a high-performance open-source super-resolution microscopy toolbox

**DOI:** 10.1101/432674

**Authors:** Romain F. Laine, Kalina L. Tosheva, Nils Gustafsson, Robert D. M. Gray, Pedro Almada, David Albrecht, Gabriel T. Risa, Fredrik Hurtig, Ann-Christin Lindås, Buzz Baum, Jason Mercer, Christophe Leterrier, Pedro M. Pereira, Siân Culley, Ricardo Henriques

## Abstract

Super-resolution microscopy has become essential for the study of nanoscale biological processes. This type of imaging often requires the use of specialised image analysis tools to process a large volume of recorded data and extract quantitative information. In recent years, our team has built an open-source image analysis framework for super-resolution microscopy designed to combine high performance and ease of use. We named it NanoJ - a reference to the popular ImageJ software it was developed for. In this paper, we highlight the current capabilities of NanoJ for several essential processing steps: spatio-temporal alignment of raw data (NanoJ-Core), super-resolution image reconstruction (NanoJ-SRRF), image quality assessment (NanoJ-SQUIRREL), structural modelling (NanoJ-VirusMapper) and control of the sample environment (NanoJ-Fluidics). We expect to expand NanoJ in the future through the development of new tools designed to improve quantitative data analysis and measure the reliability of fluorescent microscopy studies.

## Introduction

Fluorescence microscopy has been ubiquitously used in biological studies since its invention in the 20^th^ century. It can reveal subcellular structures and interactions between specifically labelled molecules and allows the quantification of their dynamic behaviour in living cells (1). Extraction of this biologically relevant quantitative information from fluorescence microscopy data typically requires digital image processing and analysis (2). In recent years, Super-Resolution Microscopy (SRM) techniques have extended the spatial resolving power of fluorescence microscopy beyond the diffraction limit (3–5). Most SRM techniques use large quantities of raw data, often reaching several gigabytes to generate a single super-resolution image, thus requiring specialised high-performance image analysis tools. Several SRM image processing packages are available, such as ThunderSTORM (6), LAMA (7) and SIMcheck (8) but each of these is focused on a specific type of SRM modality.

Here, we present NanoJ, a highly versatile set of image acquisition and analysis methods developed to improve the reliability and quantifiability of microscopy experiments, with a particular focus on the demands of live-cell SRM. NanoJ is available as a series of ImageJ-based plugins which can be used independently or concomitantly. NanoJ (Fig. 1) is comprised of the following modules: **NanoJ-Core** - general image correction tools including drift correction and channel registration, both based on cross-correlation analysis; **NanoJ-SRRF** - an analytical approach capable of extracting super-resolution data from a short sequence of diffraction-limited images, which can be acquired using most microscopes (9,10); **NanoJ-SQUIRREL** - an algorithm to evaluate resolution and the presence of artefacts in super-resolution images (11); **NanoJ-VirusMapper** - a single-particle analysis method to generate nanoscale models of biological structures such as viruses (12–14); **NanoJ-Fluidics** - a hardware and software framework to control fluidics devices, enabling automation of multiplexed experiments (15). Thus, the NanoJ framework is capable of solving common imaging problems with broad biological applications and is compatible with a multitude of fluorescence microscope setups and experimental protocols.

**Fig. 1.**
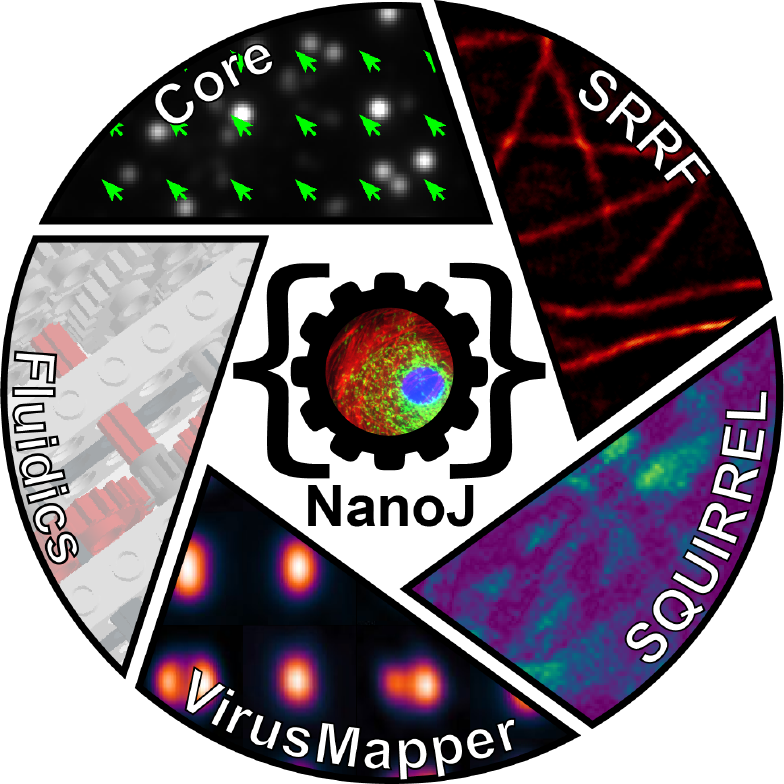
NanoJ framework. Currently NanoJ consists of 5 modules dedicated to super-resolution imaging and analysis.

## The NanoJ framework

NanoJ has been designed to integrate with the popular ImageJ or Fiji image analysis software (16, 17) and is easily installed as a standard set of plugins. NanoJ is also fully open-source and user-friendly. The graphical user interfaces (GUIs) are straightforward to use and its routines can be easily integrated within larger image analysis pipelines through the ImageJ macro language.

NanoJ is designed to be an accessible tool for both non-expert users and developers. Each of its modules possess their own separate manual and documentation. NanoJ is implemented in both Java (https://www.java.com/) and OpenCL (https://www.khronos.org/opencl), the latter language being used for high-performance analysis of image data through the use of Graphical Processing Units (GPUs). To date, it encompasses four Java ARchive (JAR) packages (NanoJ-SRRF, NanoJ-SQUIRREL, NanoJ-VirusMapper, NanoJ-Fluidics) that all depend on a central package (NanoJ-Core). The core package hosts the libraries that enable high-performance GPU-based computing analysis and a set of basic image analysis helper methods. The modular nature of NanoJ means that its components can be updated independently and the framework can be easily extended by appending new analytic packages.

## NanoJ-Core: Drift Correction

Sample drift commonly occurs during the acquisition of SRM data, often as a result of gradual changes in the temperature of microscope components. Drift introduces motion blur artefacts and thereby a loss of resolution. While most modern microscopes have an active focus-lock device that stabilises the motion of the sample in the axial direction (minimising focal drift), the sample will still be prone to lateral movement (Fig. 2a). However, in the case where the raw data is made up of a sequence of consecutive frames acquired rapidly, as is common in SRM methods such as Single Molecule Localization Microscopy (SMLM) (3, 4) or fluctuation-based approaches (9, 18, 19), this lateral drift can be estimated (Fig. 2b) and analytically corrected for each time frame via post-processing (Fig. 2c-d). NanoJ breaks the task of drift correction into two distinct parts: estimation, followed by translation. As a first step, NanoJ-Core estimates the linear drift between two images by calculating their cross-correlation matrix (CCM) (Fig. 2b). The location of the peak intensity in the CCM determines the linear shift between the two images, and precise sub-pixel accuracy is achieved by up-scaling the CCM using a bicubic spline interpolation. Depending on the type of acquisition, the reference frame can either be the first frame of the raw data or the immediately preceding frame. Fig. 2d shows the drift in a 100-frame dataset as measured with respect to the first frame. Once drift is estimated, the dataset can be directly corrected by analytically translating each individual frame using a bicubic spline interpolation (Fig. 2c). The interpolation process will, however, change the noise properties of the resulting dataset (20), which can have an influence on further analysis requiring specific assumptions in noise properties such as Maximum Likelihood Estimation (MLE) single-molecule localization analyses.

**Fig. 2.**
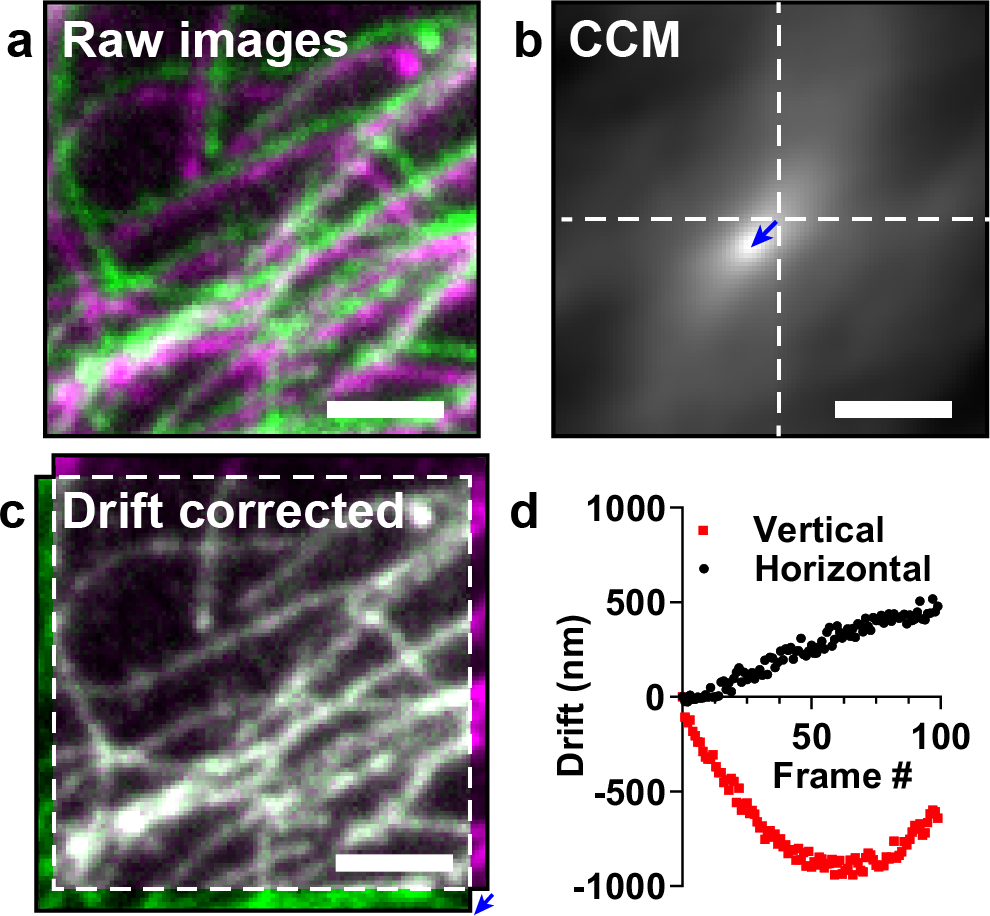
Drift correction with NanoJ-Core. **a)** Composite image of two frames from a time-lapse dataset of the same field-of-view. An artificially large drift was applied computationally in order to make it visible for figure rendering. **b)** Cross-correlation map (CCM) between the two frames shown in a). The vector position of the maximum indicates the linear shift between the two frames. **c)** Overlay of the two frames after drift correction using NanoJ-Core. **d)** Vertical and horizontal drift curves obtained using NanoJ-Core from the 100-frame raw data. The two images shown in a) correspond to the frames 0 (green) and 97 (magenta) of the raw data.

For the specific case of SMLM datasets with sparse blinking, there will only be a weak correlation across frames, as there is little observable structure conserved between consecutive time points. One common strategy to alleviate this low correlation is to add fiduciary landmarks to the sample, such as static fluorescent beads. As an option NanoJ-Core can temporally bin images within the dataset, thus increasing the correlation between frames and allowing their shift to be more accurately estimated (21).

Drift estimation in NanoJ differs from strategies applied by other SRM algorithms, such as ThunderSTORM (6), by analysing unprocessed raw data instead of post-processed super-resolution reconstructions. This allows the estimation to be decoupled from the super-resolution image reconstruction algorithm and hence drift-corrected raw data can be analysed using a wider range of methods including SRRF and SOFI(18). Furthermore, NanoJ-SRRF (as described in a following section) can import the drift curve (Fig. 2d) created by NanoJ-Core Drift Correction and use this information directly during analysis without the need to pre-translate each frame in the raw dataset.

## NanoJ-Core: Channel registration

In multicolour fluo-rescence microscopy, images acquired in different spectral channels are often misaligned as a result of chromatic aberrations in the optical path and the use of different filter sets for each colour. This misalignment is frequently ignored in conventional microscopy as it typically occurs on a scale smaller than the diffraction limit. However, this effect becomes non-negligible in the context of SRM (22). Channel registration is therefore essential for multicolour SRM studies quantifying colocalization or interactions between different structures (23–25). Advanced imaging research groups tend to develop their own analysis scripts for their particular applications, and thus no user-friendly tools are readily available. Therefore, NanoJ-Core Channel registration offers a unique tool to perform this registration easily and robustly on a wide variety of datasets.

The shift between different spectral channels is usually inhomogeneous across a field of view, which prevents the use of typical CCM-based approaches such as the one described for NanoJ-Core Drift Correction. In order to characterise the spectral misalignment across a field of view, it is necessary to image a sample where the same structure can be observed across all spectral channels of interest. Furthermore, the sample should contain structures occupying the whole field of view. A typical sample for this characterisation is a coverslip coated with a large number of beads labelled with multiple fluorescent dyes (26). Following multicolour imaging of this sample, NanoJ-Core can be used to calculate a non-linear two-dimensional spatial transform describing the mis-alignment for each channel relative to a reference channel. Given that chromatic misalignment is a fixed property of an optical system, NanoJ-Core can then apply these transforms to realign other multicolour datasets acquired using the same optical path (Fig. 3) (27, 28).

**Fig. 3.**
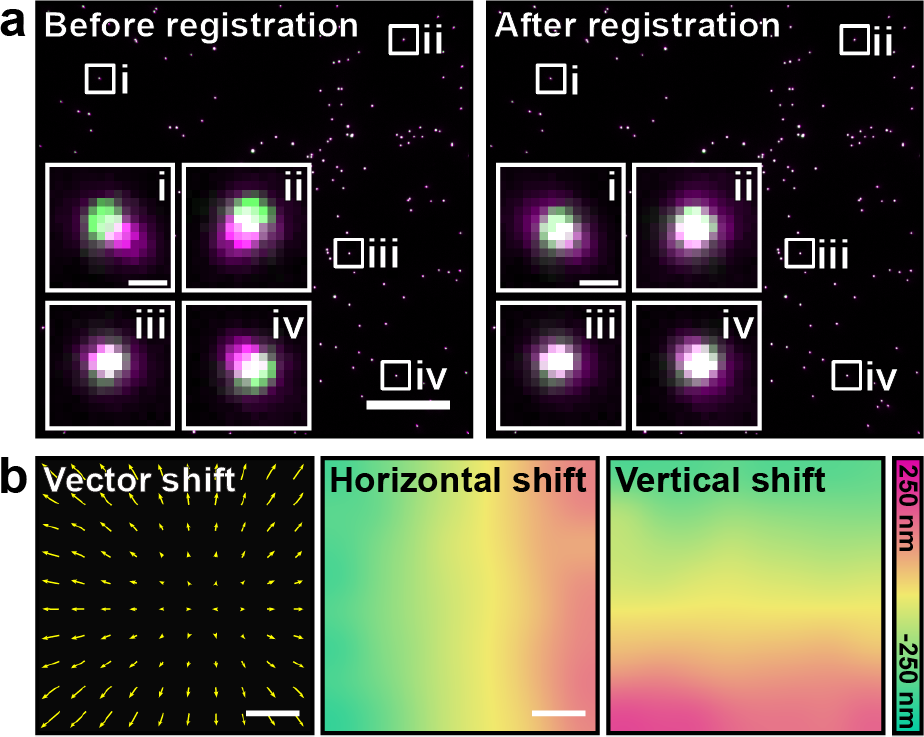
Multi-colour channel registration with NanoJ-Core. **a)** Composite image of multi-colour TetraSpeck™ beads imaged in two different channels (“GFP-channel” indicated in green and “mCherry-channel” in magenta), prior to (left) and after (right) channel registration using NanoJ-Core. Insets - individual beads from indicated locations. Scale bars: 25 μm, insets: 0.5 μm. **b)** Vectorial representation of the shift between the two channels (left, displacement vector length 50 times larger for representation purposes), horizontal (middle) and vertical (right) shift maps obtained and applied to the data shown in a). Scale bars: 25 μm.

To generate these non-linear misalignment fields NanoJ-Core first calculates local, linear misalignments between channels (Fig. 3a). This is achieved by dividing the image into small areas (“blocks”). For each block, the shift is calculated by finding the cross-correlation peak position as shown in Fig. 2a, b. These local shift values are then interpolated across the whole field-of-view using an inverse distance weighting interpolation (29). This generates two smooth shift maps describing the misalignments in the horizontal and vertical directions (Fig. 3b). For a given channel, each pixel value within the horizontal/vertical shift map indicates the horizontal/vertical displacement that needs to be applied to that pixel to align it with the reference channel. NanoJ-Core can then be used to apply these maps to align channels in the dataset of interest (Fig. 3), provided they have been acquired using the same optical configuration.

NanoJ-Core performs the channel registration by creating a new image representing each channel, where the intensity value for each pixel coordinate corresponds to the intensity value from the original image at the equivalent coordinate corrected for local shift. For cases in which these coordinates are not discrete (sub-pixel shift), a bicubic spline interpolation is used to recover pixel values in continuous space. Because the shift map can be extrapolated to continuous space, the registration procedure obtained from diffraction-limited images can also easily be used to correct super-resolved images obtained using the same optical configuration.

## NanoJ-SRRF: Live-Cell Super-Resolution Imaging

As part of the NanoJ framework, we include our recently developed SRM reconstruction algorithm Super-Resolution Radial Fluctuations (SRRF), which is able to extract sub-diffraction information from a short burst of images acquired at high-speed with modern fluorescence microscopes (9, 10). SRRF is a purely analytical approach. It alleviates the need to use toxic photoswitching-inducing buffers (30), specialised fluorophores (31, 32), damaging high-intensity illumination (33) or specialised equipment (5, 34) when compared to other SRM methods (3–5, 34).

SRRF is based on similar principles to SMLM, with the key difference that it does not rely on the detection of spatio-temporally isolated fluorophores. Instead, SRRF generates a magnified pixel grid where each pixel value relates to the probability of fluorophores existing in that corresponding region of space. To do this, SRRF calculates the local radial symmetry (‘radiality’) in each pixel of the magnified image using local intensity gradient information. The obtained radiality value will be high when a point-spread-function (PSF) profile transiently becomes dominant, highlighting the presence of a fluorescent molecule at that location. Furthermore, the fluctuations of radiality values follow the underlying natural intensity fluctuations of fluorophores, which have a distinct temporal signature to that of noise (18). Therefore, a temporal correlation of the radiality at each pixel can be projected into a final image, where the structures of interest will be better resolved.

Thanks to its low-illumination requirement, SRRF is highly suited to enable live-cell SRM and allows for long SRM time-lapse acquisitions (10). Fig. 4 shows a typical live-cell SRRF acquisition of a Cos7 cell expressing UtrCH-GFP (a probe for actin filaments) imaged at 33.3 Hz. The dataset shown is part of a longer > 30 min time-course dataset (Fig. S2). SRRF allows the observation of protein dynamics (e.g. actin) during long periods with no perceivable phototoxicity.

**Fig. 4.**
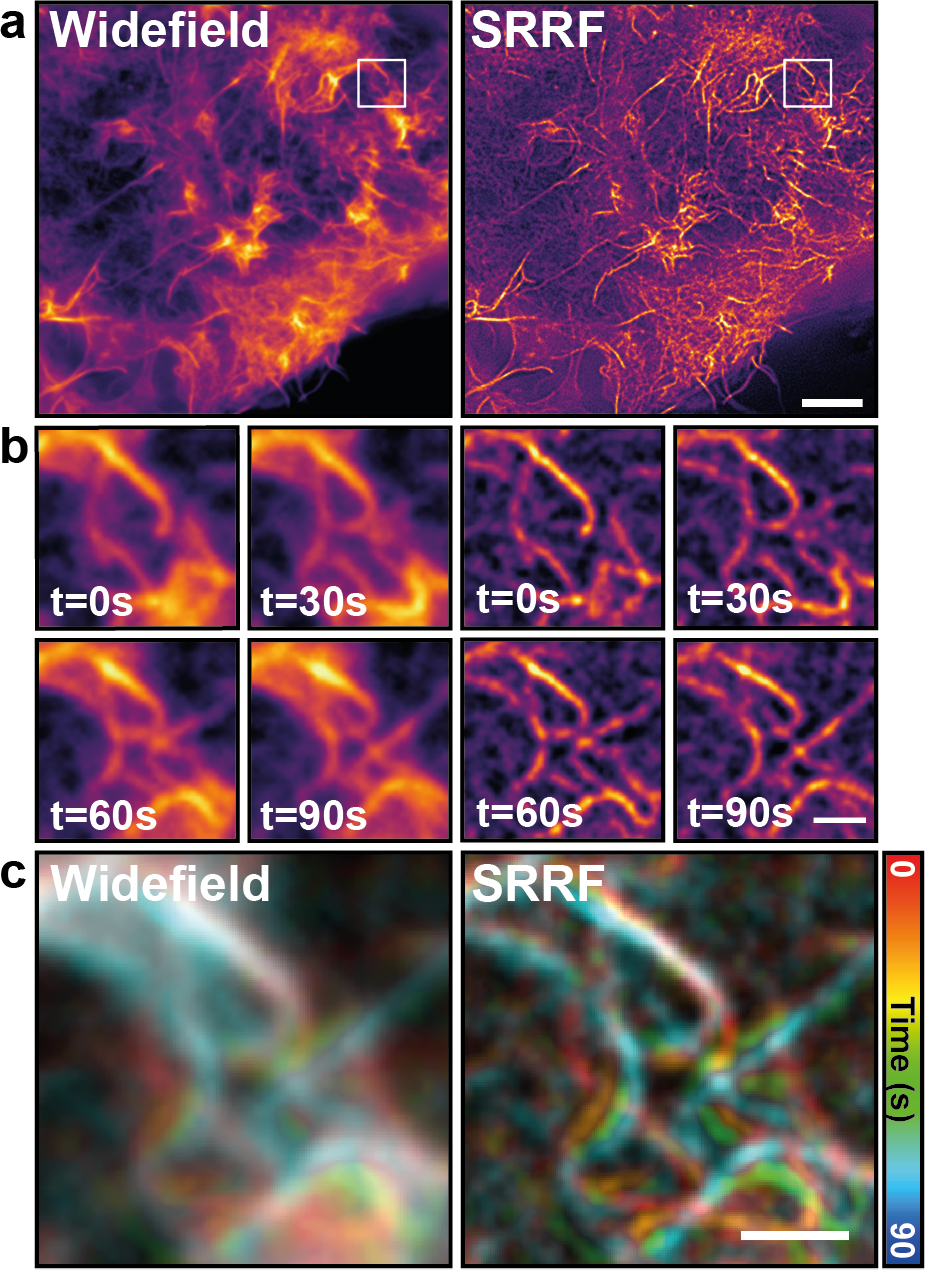
Live-cell super-resolution microscopy with NanoJ-SRRF. **a)** Comparison of widefield (left) and SRRF reconstruction (right) obtained from a Cos7 cell expressing UtrCH-GFP to label actin filaments. Scale bar: 5 μm. **b)** Time-course of the inset shown in a), obtained from a continuous imaging at 30 ms exposure (33.3 Hz) and displayed every 30 s. Scale bar: 1 μm. **c)** Colour-coded time course dataset from b). Scale bar: 1 μm.

NanoJ-SRRF, the software implementation of the SRRF algorithm, uses GPU computing whenever possible to accelerate radial symmetry calculations. This is achieved by implementing part of its analytic engine in OpenCL, with a fallback of execution to the CPU if a compatible graphics card is not detected.

## NanoJ-SQUIRREL: Estimating Image Quality

SRM techniques are more complex than conventional diffraction-limited microscopy. This complexity arises from the sample preparation requirements (SMLM), the microscope hardware (stimulated emission depletion microscopy or STED, Structured Illumination Microscopy or SIM) and the post-acquisition image processing (SMLM, SIM). Unsuitable choices in any of these underlying variables can lead to artefacts within the final image and the potential for false conclusions to be drawn. NanoJ-SQUIRREL (Super-resolution Quantitative Image Rating and Reporting of Error Locations) is an algorithm that highlights the presence of artefacts in super-resolution images. It does so by calculating quantitative maps showing both local SRM image quality and resolution (11) and can thus be used to optimise acquisition workflows (10).

The central concept of SQUIRREL is that a diffraction-limited image and the corresponding SRM image of the same region should contain the same underlying structure, just at different resolutions. Thus, the diffraction-limited image can be treated as a high-confidence benchmark against which the SRM image can be compared. The SQUIRREL image quality assessment algorithm formalises this comparison analytically.

NanoJ-SQUIRREL quality assessment requires the user to provide an acquired diffraction-limited image and the SRM image of the same structure (Fig. 5a). As both images represent the underlying fluorophore distribution but at different resolutions, there exists a blurring function that can convert the SRM image into its diffraction-limited equivalent. SQUIRREL calculates this blurring function and applies it to the SRM image. The blurred image is then compared against the diffraction-limited reference. This generates three quality indicators. An error map is produced that maps the discrepancy between the blurred SRM image and the reference image at every pixel (Fig. 5b). This highlights regions where the super-resolution image is inconsistent with the reference image. For example, the three insets in Fig. 5b show typical SMLM artefacts including incomplete structures and inter-structure mislocalization. Two global quality metrics are also generated for each image: the RSP (Resolution Scaled Pearson’s Correlation Coefficient) and the RSE (Resolution Scaled Error). The RSP can take a value in the interval [−1,1] and describes the structural agreement between the blurred super-resolution and reference images. Here, higher values indicate better agreement, with an RSP of 1 indicating a perfect structural match. The RSE describes the mean intensity mismatch between the blurred super-resolution and reference images; in this case lower values represent better agreement with a value of 0 indicating a perfect intensity match.

## NanoJ-SQUIRREL: Estimating Image Resolution

The purpose of SRM is to resolve finer structural detail than is achievable with conventional diffraction-limited microscopy. It is therefore useful to have an objective measurement of resolution within a super-resolution image, for example to enable meaningful analysis and structural hypotheses that stay in line with the actual precision of the data. The current standard for measuring image resolution in SMLM images is Fourier Ring Correlation (FRC) (35). This method involves comparing two independently acquired super-resolution images of the same field-of-view so that they only differ by their noise components. For SMLM datasets the two SRM images are usually obtained by splitting localizations from odd and even frames. The correlation between these two SRM images is measured at different frequencies in Fourier space; the frequency at which this correlation drops below a set threshold indicates the resolution of the image.

FRC has been previously implemented in ImageJ (35), but only gives a single resolution measurement for the entire field of view. However, resolution is not necessarily homogeneous across the SRM image (Fig. 5c). This is particularly true for SMLM methods as localization accuracy depends strongly on labelling density and laser illumination intensity, which can both vary considerably within a single field of view. Furthermore, FRC can generate biased measurements for certain fluorophore distributions such as point-like patterns. Therefore, an additional feature of the NanoJ-SQUIRREL plugin is local mapping of FRC resolution across an image. To do this, the user provides an image stack comprising two independent renderings of the same dataset (e.g. through the odd/even frames splitting as described above). The images are then spatially split into equal-sized blocks and FRC analysis is run locally on each block. For blocks where there is insufficient correlation to generate an FRC resolution value, a resolution value is interpolated from neighbouring blocks. Fig. 5c shows the FRC map obtained from the SRM image shown in (a). This map highlights that the resolution in this image varies between 85 and 36 nm.

**Fig. 5.**
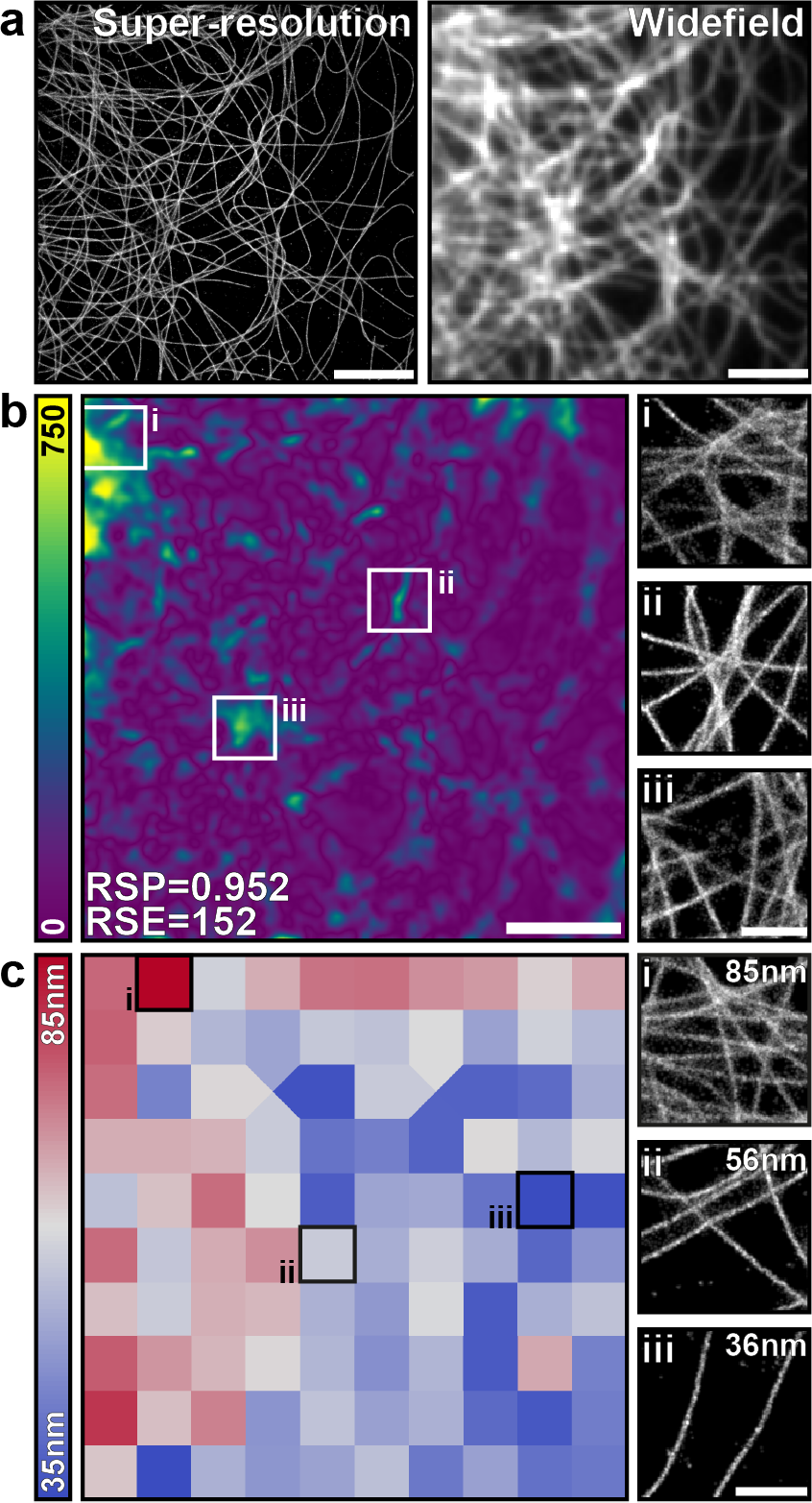
Quality assessment and resolution mapping with NanoJ-SQUIRREL. **a)** A super-resolution rendering (left) and acquired widefield image (right) of fixed microtubules labelled with Alexa Fluor-647. **b)** Left: SQUIRREL error map highlighting discrepancies between the super-resolution and diffraction-limited images in (a). Right: Magnified insets of super-resolution rendering at indicated positions on error map. **c)** Left: SQUIRREL resolution map of the super-resolution image in (a). Right: Magnified insets of super-resolution rendering for indicated resolution blocks. Whole image scale bars= 5μm, inset scale bars = 1μm

It is important to note that high resolution (that is, a low FRC value) does not imply that the super-resolution image has depicted structures correctly; it only means that there is low variation in the locations of the fluorophores between the two rendered images. Therefore, the error mapping functionality within NanoJ-SQUIRREL can complement FRC-mapping in order to obtain a more complete perspective on super-resolution image quality.

## NanoJ-VirusMapper: Structural Mapping and Modelling

As part of the NanoJ framework, we include a unique single-particle analysis (SPA) tool called NanoJ-VirusMapper. It is the first open-source, freely available algorithm for unbiased, high-throughput SPA of fluorescence imaging and allows the structural modelling of viruses and other macromolecular complexes (12–14). The principle of SPA is to image many identical copies of a structure, independently of its orientation, and align and combine them to build an averaged structural map of the underlying structure with high signal-to-noise ratio (36–40).

**Fig. 6.**
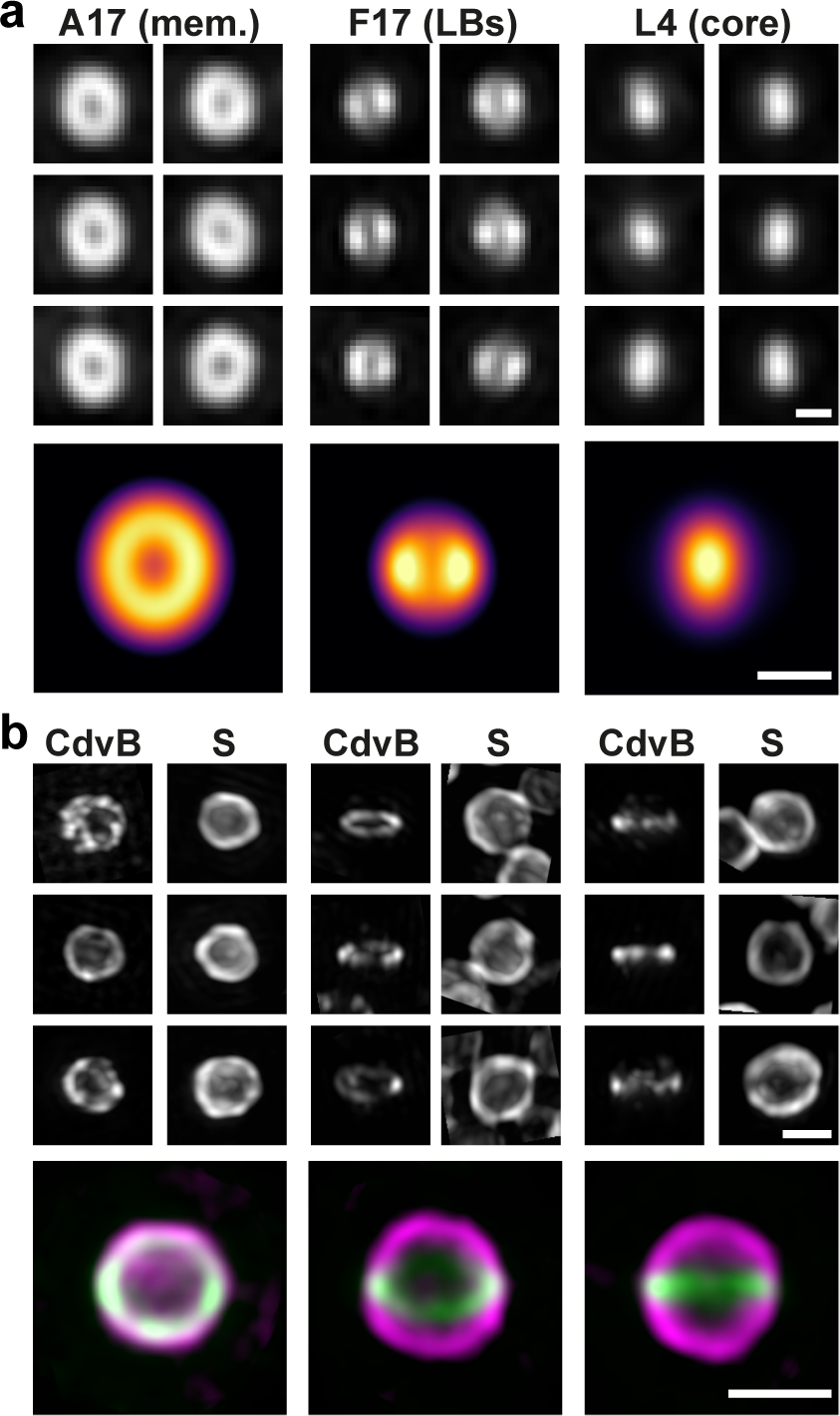
Quantitative SPA-based modelling with NanoJ-VirusMapper. **a)** Top: Aligned SIM images of individual vaccinia particles labelled for L4 (core), F17 (lateral bodies, LBs) and A17 (membrane, mem.). Bottom: VirusMapper models of the three channels. Scale bars: 200nm. **b)** Top: Aligned SIM images of individual *Sulfolobus acidocaldarius* cells labelled for the S-layer (S) and the archaeal ESCRT-III homolog CdvB (CdvB). Bottom: VirusMapper models of three different orientations of the cells; magenta-S-layer, green-CdvB. Scale bars: 1μm.

The SPA implementation of VirusMapper facilitates automatic processing of multiple images to detect, segment, align, classify and average thousands of individual structures. It is entirely general, assuming no underlying symmetry or other properties of the imaged structure. Here, we illustrate this with models of three distinct vaccinia virus substructures (41): core, lateral bodies (LBs) and membrane. All three sub-structures were labelled on viral particles and imaged with SIM (Fig. 6a, top). VirusMapper can explicitly incorporate information from multiple fluorescence channels to enable multi-component modelling. Here, the images of the LBs and the core were used to find the orientations of each particle in the dataset. This allowed us to create simultaneous models of the three components of the virus (Fig. 6a, bottom). VirusMapper can also be used to separately model different 3D orientations of a structure. Here, we highlight this capability on a study of the division ring in the thermoacidophilic archaeon *Sulfolobus acidocaldarius*, strain DSM639 (Fig. 6b). *Sulfolobus* cells were labelled for both their outer S-layer and the division ring, as marked by the archaeal ESCRT-III homolog CdvB. The images of the division ring were used to identify different orientations of the cells (Fig. 6b) and separate templates were created using VirusMapper. Parallel models were then generated of the division ring and the S-layer using these templates for each orientation. The models obtained from the two channels can then be overlayed (Fig. 6b, bottom).

VirusMapper is compatible with any fluorescence microscopy method and has been demonstrated with SIM, STED microscopy (12) and SMLM (Fig. S1). The general approach of the method enables it to even be applied to correlative combinations of methods (Fig. S1).

## NanoJ-Fluidics: Sample Liquid Exchange

NanoJ-Fluidics is a hardware and software framework for precise and accurate automated liquids exchange (15). It was developed to enable automation of sample treatment and labelling of live or fixed specimens directly on the microscope stage (15, 42). The NanoJ-Fluidics hardware component is composed of customisable, low-cost and robust LEGO^®^ syringe pumps and a liquid removal peristaltic pump, all controlled by simple Arduino^®^ electronics. It is compatible with off-the-shelf imaging chambers, without the need for any micro-fabrication. Its control software (the NanoJ-Fluidics module) is ImageJ-based and can be fully integrated with microscopy acquisition software. We have demonstrated the applicability of NanoJ-Fluidics in multiple experimental contexts, including *in-situ* correlative live-to-fixed super-resolution imaging, multimodal super-resolution imaging and event-driven fixation (15). The approach can also be easily extended to protocol optimisation (e.g. titrating antibody concentrations or adjusting imaging buffer composition) or liquid exchange protocols integrated with the imaging (e.g. drug delivery or automated event-driven fixation).

Here, we demonstrate NanoJ-Fluidics by acquiring a high-quality multicolour SMLM dataset, where STORM and DNA-PAINT (43) imaging strategies are combined into a single workflow (Fig. 7a). This approach is particularly suited to multi-target imaging (31), which is difficult to achieve with standard sample preparation techniques due to the low number of suitable fluorophores available for SMLM microscopy. With NanoJ-Fluidics, we can seamlessly perform all labelling steps in an automated and reliable manner directly on the microscope stage. We showcase this using a four-channel acquisition of actin with STORM, and mitochondria, vimentin and clathrin with DNA-PAINT (Fig. 7b).

**Fig. 7.**
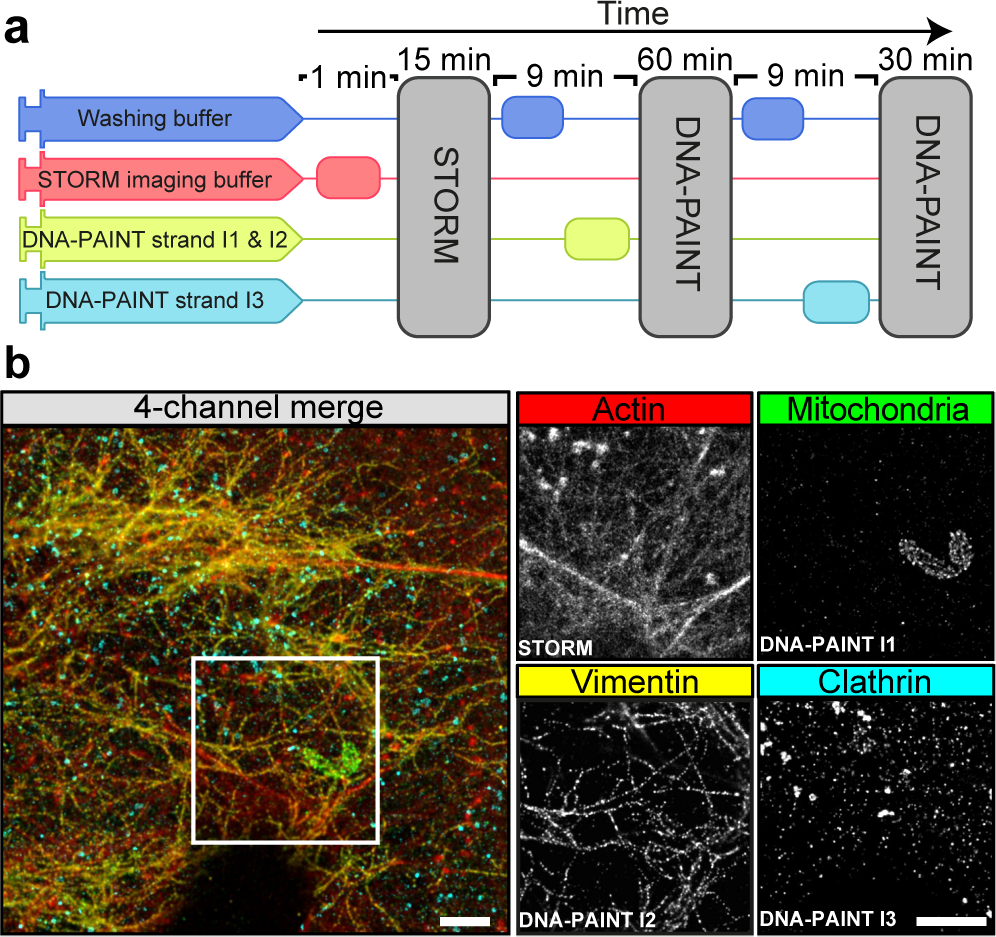
Automated DNA-PAINT and STORM imaging. **a)**NanoJ-Fluidics workflow used for multi-color STORM and DNA-PAINT imaging. **b)** Left: 4-channel merge of STORM and DNA-PAINT with actin (red, STORM), mitochondria (green, DNA-PAINT I1 strand), vimentin (yellow, DNA-PAINT I2 strand) and clathrin (cyan, DNA-PAINT I3 strand). Right: Single-channel images from the inset. Scale bars: 2 μm.

NanoJ-Fluidics’ highly customisable nature has already spawned several alternative designs from the community (https://github.com/HenriquesLab/NanoJ-Fluidics/wiki), which are in constant development. NanoJ-Fluidics makes imaging protocol automation readily available to researchers, hence improving not only the reliability and repeatability of the protocols, but also the range of protocols that are achievable.

## Discussion and Future Perspectives

The NanoJ frame-work provides a unique and comprehensive set of tools to support fluorescence imaging, from data acquisition and protocol optimisation to structural quantification. It offers powerful and user-friendly solutions to common pitfalls of image analysis such as drift correction and channel registration (NanoJ-Core). It extends the tools available for SRM to expert and novice users alike and aligns with the general aim of the community to extend SRM to live-cell applications, offering the first open-source analytical approach for live-cell SRM imaging (NanoJ-SRRF). NanoJ offers a way to correlate dynamic live-cell data with high-resolution fixed-cell data in an automated and highly reproducible manner (NanoJ-Fluidics). It incorporates an analytical method for the generation of accurate, high-content, molecular specific models (NanoJ-VirusMapper). Finally, the NanoJ framework provides tools to quantitatively assess SRM image quality by spatially mapping unbiased quality metrics, artefacts (errors) and FRC resolution (NanoJ-SQUIRREL), with the aim to improve standards in assessing and reporting microscopy data. The NanoJ framework gives access user-friendly yet robust analytical methods in tandem with new approaches to perform live-cell SRM experiments. A typical SRM live-cell imaging experiment using the NanoJ framework would entail acquiring a SRRF-compatible raw data set in any modern fluorescence microscope; performing drift correction and channel registration; followed by SRRF analysis for SRM image reconstruction; and finally, quality check using SQUIRREL. Used concomitantly in this manner, these tools can be used to address a large variety of biological questions and can be seamlessly repeated for imaging protocol optimisation and complemented using NanoJ-VirusMapper and/or NanoJ-Fluidics. Thus, the NanoJ-enabled scientific pipeline allows any user, regardless of the experience level, to seamlessly obtain SRM quantitative data of the highest scientific quality. The NanoJ framework was developed to facilitate access and expand the options of research groups using fluorescence microscopy. Hence, speed, reliability, performance and cross-compatibility are of paramount importance. In this context, NanoJ has been designed for high-performance image analysis, using GPU computing, ensuring the quick processing of large data volumes (44–48). Further, NanoJ’s modular and open-source nature and its integration within ImageJ allow it to be used in conjunction with other analysis software packages (49–52). We are continuously supporting, adapting and expanding the framework to include new approaches, such as 3D imaging.

We expect NanoJ to set the standard for useful, open-source, high performance methods for the whole microscopy community.

## Software and Hardware Availability

NanoJ follows open-source software and hardware standards. Each of its modules can be installed by enabling the corresponding code repository in Fiji or by following the instructions on the corresponding websites:

- https://github.com/HenriquesLab/NanoJ-Core
- https://github.com/HenriquesLab/NanoJ-SRRF
- https://bitbucket.org/rhenriqueslab/NanoJ-SQUIRREL
- https://bitbucket.org/rhenriqueslab/NanoJ-VirusMapper
- https://github.com/HenriquesLab/NanoJ-Fluidics

## ACKNOWLEDGEMENTS

First and foremost, we would like to thank the ImageJ and Fiji open-source development community. Their astonishing work has been the inspiration to the creation of NanoJ. There has been a large number of beta-testers and NanoJ users whose feedback helped us to continually update and improve its code, to the best of our capacity we keep an updated list of significant contributors in the website corresponding to each NanoJ module. We thank Prof. Ralf Jungmann at Max Planck Institute of Biochemistry Munich for reagents and advice. This work was funded by grants from the UK Biotechnology and Biological Sciences Research Council (BB/M022374/1; BB/P027431/1; BB/R000697/1; BB/S507532/1) (R.H., P.M.P. and R.F.L.), the UK Medical Research Council (MR/K015826/1) (R.H.), the Wellcome Trust (203276/Z/16/Z) (S.C., R.H and B.B.), Core funding to the MRC Laboratory for Molecular Cell Biology, University College London (MC_UU12018/7) (J.M.), the European Research Council (649101–UbiProPox) (J.M.) and the Centre National de la Recherche Scientifique (CNRS ATIP-AVENIR program AO2016) (C.L.). N.G. and R.D.M.G funded by the Engineering and Physical Sciences Research Council (EP/L504889/1). P.A. was supported by a PhD fellowship from the UK’s Biotechnology and Biological Sciences Research Council. Research by B.B. was supported by UCL, Cancer Research UK (C1529/A17343), BBSRC (BB/P001440/1) and MRC (MC_CF12266). K.L.T. and G.T.R. are supported by a 4-year MRC Research Studentship. D.A. is presently a Marie Curie fellow (Marie Sklodowska-Curie 750673). A.C.L. AND F.H. were supported by the Swedish Research Council, grant 621-2013-4685.

## AUTHOR CONTRIBUTIONS

These contributions follow the Contributor Roles Taxonomy guidelines: https://casrai.org/credit/. Conceptualization: K.L.T, R.F.L., N.G., R.D.M.G, P.A., D.A., G.T.R., B.B., J.M., C.L., P.M.P., S.C., R.H.; Data curation: K.L.T, R.F.L., R.D.M.G, G.T.R., J.M., C.L., P.M.P., S.C., R.H.; Formal analysis: K.L.T, R.F.L., R.D.M.G, C.L., S.C.; Funding acquisition: A.C.L, B.B., C.L., R.H.; Investigation: K.L.T, R.F.L., R.D.M.G, G.T.R., C.L., P.M.P., S.C.; Methodology: R.F.L., N.G., R.D.M.G, P.A., J.M., C.L., P.M.P., S.C., R.H.; Project administration: R.F.L., B.B., J.M., C.L., P.M.P., S.C., R.H.; Resources: K.L.T, R.D.M.G, D.A., G.T.R., F.H., A.C.L., B.B., J.M., C.L., P.M.P., S.C.; Software: R.F.L., N.G., R.D.M.G, P.A., C.L., P.M.P., S.C., R.H.; Supervision: R.F.L., A.L., B.B., J.M., C.L., P.M.P., S.C., R.H.; Validation: K.L.T, R.F.L., N.G., R.D.M.G, P.A., D.A., C.L., P.M.P., S.C.; Visualization: K.L.T, R.F.L., R.D.M.G, D.A., C.L., P.M.P., S.C., R.H.; Writing – original draft: K.L.T, R.F.L., R.D.M.G, D.A., C.L., P.M.P., S.C., R.H.; Writing – review & editing: all authors.

## Supplementary Note 1: Drift correction

FluoCells slide #2 (Thermo Fisher Scientific) were imaged on a Ti2 microscope (Nikon) (Fig. 2) using a 100x TIRF objective (CFIApochromat TIRF 100XC Oil, Nikon) onto a Prime 95B camera (Photometrics) in the GFP channel (tubulin), using an LED illumination source. 100 frames were acquired at 40 Hz and an artificially large drift was applied digitally.

## Supplementary Note 2: Channel alignment

Multicolor beads (TetraSpeck™ Microspheres, 0.1 μm, fluorescent in blue/green/orange/dark red, Thermo Fisher Scientific) were prepared on a #1.5 cover slip and embedded in water before being sealed with nail polish. The beads were then imaged on a Ti2 microscope (Nikon) (Fig. 3) using a 100x TIRF objective (CFI Apochromat TIRF 100XC Oil, Nikon) onto a Prime 95B camera (Photometrics), in both GFP and mCherry channels, using an LED illumination source.

## Supplementary Note 3: SRRF

### A. Cell lines

Cos7 cells were cultured in phenol red-free Dulbecco’s modified Eagle’s medium (DMEM; Thermo Fisher Scientific) supplemented with 10% (v/v) fetal bovine serum (FBS; Gibco), 1% (v/v) penicillin/streptomycin (Thermo Fisher Scientific) and 2 mM L-alanyl-L-glutamine (GlutaMAX™, Thermo Fisher Scientific) at 37 °C in a 5% CO_2_ incubator.

### B. Sample preparation

For live-cell imaging, Cos7 cells were seeded on 25mm-diameter #1.5 coverslips (Marienfeld) at a density of 0.3 −0.9×10^5^ cells/cm^2^. One day after splitting, cells were transfected with a plasmid encoding the calponin homology domain of utrophin fused to GFP (GFP-UtrCH) (1) using Lipofectamin 2000 (Thermo Fisher Scientific) according to the manufacturer’s recommendations. Cells were imaged 1–4 days post transfection in culture medium.

### C. Imaging

LED-illumination widefield imaging of GFP-UtrCH in live Cos7 cells (Fig. 4) was performed in at 37 °C and 5% CO_2_ on a N-STORM microscope (Nikon). A 100x TIRF objective (Apochromat 100x/1.49 Oil, Nikon) with additional 1.5x magnification was used to collect fluorescence onto an EMCCD camera (iXon Ultra 897, Andor), yielding a pixel size of 107 nm. Frames were acquired for 30 min with 30 ms exposure and 490 nm LED illumination at 5% of maximum output.

### D. SRRF reconstruction

The dataset was reconstructed using NanoJ-SRRF with the following parameters: magnification: 5, radius: 0.5, number of axes: 6, number of frames per SRRF reconstruction: 100, temporal analysis: Temporal Radiality Average (TRA).

## Supplementary Note 4: SQUIRREL

### A. Cell lines

Cos7 cells were cultured as described in Supplementary Note 3.

### B. Sample preparation

Cos7 cells were fixed with glutaraldehyde and labelled with two monoclonal mouse anti-alpha tubulin antibodies (DM1A and B-5-1-2, both from Sigma) and a goat anti-mouse Alexa Fluor 647-conjugated secondary antibody (A21235, Thermo Fisher Scientific). Samples were mounted in Smart Buffer (Abbelight) for imaging.

### C. STORM acquisition and reconstruction

STORM imaging of Cos7 cells (Fig. 5) was performed on an N-STORM (Nikon) microscope using a 100x TIRF objective (Apochromat 100x/1.49 Oil, Nikon). Prior to STORM imaging, a reference widefield image was acquired using low intensity 642 nm laser excitation. For subsequent STORM imaging, 60,000 frames were acquired with high intensity 642 nm laser excitation at 15 ms exposure with pixel size of 160 nm and 0.1248 photons per analog-to-digital unit. Localizations were detected using the N-STORM software (Nikon), and exported as a text file to be rendered using ThunderSTORM (2).

### D. SQUIRREL analysis

The initially acquired widefield image was used as the reference and the ThunderSTORM reconstruction as the super-resolution image. All parameters in the plugin were left at their default values. For local resolution measurements, two independent super-resolution images were created using ThunderSTORM; one comprising of the localisations from odd-numbered frames, and the other of even-numbered frames. These images were concatenated to form a two slice image stack and the ‘Calculate FRC-map’ method of NanoJ-SQUIRREL was run with 10 blocks per axis.

## Supplementary Note 5: VirusMapper

### A. Vaccinia virus sample preparation

The L4-mCherry F17-EGFP virus was based on the WR strain and was described previously as WR EGFP-F17 VP8-mCherry (3). A4-EGFP was described previously as A5-EGFP (4). Virions were produced in BSC-40 cells, purified from cytoplasmic lysates and banded on a sucrose gradient. High performance coverslips (18×18 mm, #1.5H Zeiss) were washed with water, ethanol and acetone sequentially three times, then sonicated for 20 min in 1 M potassium hydroxide. The purified virus was diluted in 1 μL of 1 mM Tris buffer (pH 9) and placed onto the coverslips. After 30 min, virus solution was removed and the virus was fixed with 4% formaldehyde in PBS for 20 min. Samples were washed 3 times with PBS and blocked for 30 min with 5% BSA in PBS. A17 was stained with rabbit polyclonal anti-A17 antibody, which was a kind gift from Jacomine Krijnse-Locker (Institut Pasteur, Paris, France) and Alexa Fluor 647-conjugated secondary antibody (A21245, Thermo Fisher Scientific). For STORM imaging, A4-EGFP virus was bound to coverslips as described above. Samples were permeabilised with 0.2% Triton-X 100 for 10 min, blocked 30 min with 5% BSA in PBS and labelled with anti-L1 antibody 7D11, which was purified from a mouse hybridoma cell line kindly provided by Bernard Moss (NIH, Bethesda, MD) with permission of Alan Schmaljohn (University of Maryland, Baltimore, MD), and Alexa Fluor 647-conjugated secondary antibody (A21235, Thermo Fisher Scientific).

### B. *Sulfolobus acidocaldarius* sample preparation

*Sulfolobus acidocaldarius* strain DSM639 was grown in Brock’s medium (pH 3.2), supplemented with 0.1% NZ-amine and 0.2% sucrose at 75 ºC. Cells were fixed at OD600: 0.310 (i.e. within the limit of exponential growth) in 70% EtOH, rehydrated in PBST (0.2% v/v Tween) and stained overnight with polyclonal rabbit antibody for CdvB (5). Secondary staining was done with Alexa Fluor 647-conjugated Concanavalin A-antibody (C21421, Thermo Fischer Scientific) for labelling the S-layer and Alexa Fluor 546-conjugated secondary goat anti-rabbit antibody (A11035, Thermo Fischer Scientific) for visualising CdvB. The cells were spun down onto 2% polyethylenimine-coated coverslips before imaging.

### C. SIM imaging

SIM imaging of vaccinia virus (Fig. 6a) was performed using a 63x objective (Plan-Apochromat 63x/1.4 oil DIC M27, Zeiss) and *Sulfolobus* imaging (Fig. 6b) was performed using a 100x TIRF objective (alpha Plan-Apochromat 100x/1.46 Oil DIC M27, Zeiss), both on an Elyra PS.1 microscope (Zeiss). Images were acquired using 5 phase shifts and 3 grid rotations, with the 647 nm (32 μm grating period) and 561nm (32 μm grating period) and the 488 nm (32 μm grating period) lasers, and filter set 3 (1850–553, Zeiss). 2D-(vaccinia) and 3D-images (*Sulfolobus*) were acquired on an sCMOS camera and processed using ZEN Black software (Zeiss). Channels were aligned with the NanoJ-Core Channel Alignment Tool.

### D. SIM/STORM imaging

Correlative SIM/STORM imaging of vaccinia virus (Fig. S1) was performed on a Zeiss Elyra PS.1 inverted microscope with a 100x TIRF objective (alpha Plan-Apochromat 100× /1.46 NA oil DIC M27, Zeiss) objective with a 1.6x tube lens and an iXon 897 EMCCD camera (Andor). SIM imaging was performed as described above. STORM images were acquired at 25-30 ms exposure with 642 nm excitation at 100% laser power and a 655 nm LP filter. Activation was dynamically controlled with a 405 nm laser at 0-2% laser power. Images were processed using ThunderSTORM (2). Localisations were fitted with a maximum-likelihood estimator, lateral drift corrected by cross-correlation and localisations <20 nm apart within 2 frames merged.

### E. VirusMapper processing

Individual viral particles or archaea cells were extracted from the SIM or SIM/STORM images. Template images were generated with VirusMapper as described previously (6). 2- or 3-colour models of vaccinia virus proteins and 2-colour models of *Sulfolobus* S-layer and division ring in different orientations were then created by registration of the entire set of particles according to cross-correlation with the templates and calculation of a weighted average of a subset of particles.

## Supplementary Note 6: NanoJ-Fluidics

### A. Sample preparation

Sample were prepared as described previously (7). Briefly, Cos7 cells were grown, seeded on glass coverslips and then fixed as described in Supplementary Note 4 and were labelled with primary antibodies overnight: mouse monoclonal anti-TOM20 (612278, BD Biosciences), rabbit polyclonal anti-clathrin heavy chain (ab21679, Abcam) and chicken poly-clonal anti-vimentin (919101, BioLegend). Cells were then incubated with Exchange-PAINT secondary antibodies coupled to DNA sequences (7): goat anti-mouse I1, goat anti-chicken I2 and goat anti-rabbit I3. After rinses, the cells were incubated with 12.5 μM Atto488-conjugated phalloidin (Merck) for 90 min at room temperature and imaged within 3-4 days.

### B. NanoJ-Fluidics workflow and SMLM acquisition

Imaging of Cos7 cells (Fig. 7) was performed as described in (7). The NanoJ-Fluidics pump array was installed on an N-STORM microscope (Nikon) using a 100x TIRF objective (CFI Apochromat 100x/1.49 Oil, Nikon). First, a STORM imaging of phalloidin-Atto488 was performed using 30,000 frames at 30 ms exposure. After injection of the I1-Atto655 (0.25 nM) and I2-CY3B (2 nM) imager strands, 60,000 frames were acquired in an alternating way (30,000 frames of each channel) to image TOM20 and vimentin, respectively. After rinses, I3-Cy3B (1 nM) was injected, and 30,000 frames were acquired to image clathrin.

### C. SMLM reconstruction

Localisations were detected using the N-STORM software (Nikon), and exported as a text file before being filting (number of photons between 700 and 50,000; number of detections (after linking across frames) < 50 frames) and rendered using ThunderSTORM.

**Fig. S1.**
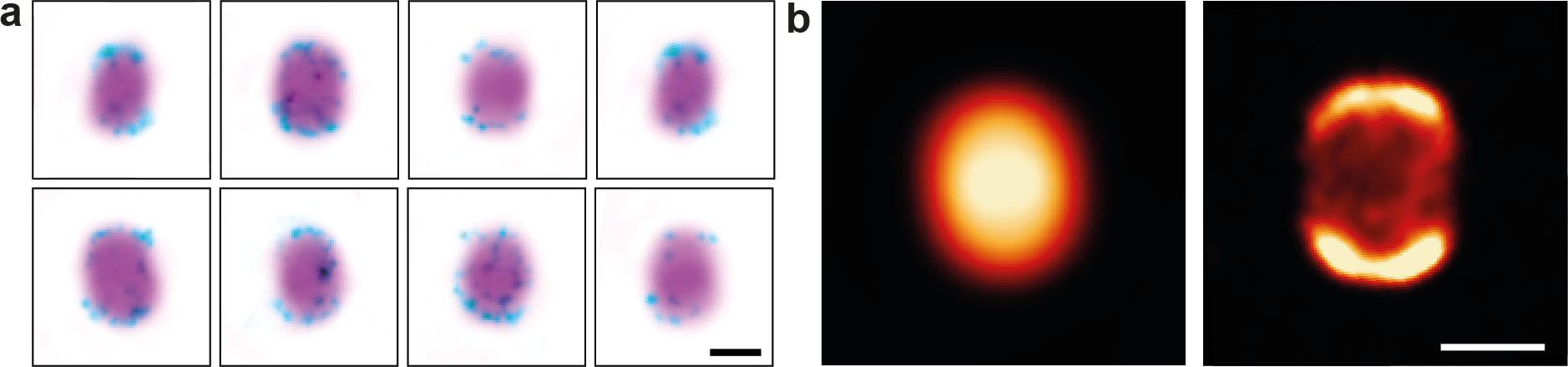
NanoJ-VirusMapper analysis of SIM/SMLM data. **a)** Example SIM/STORM images of A4-EGFP vaccinia virions labelled for the entry protein L1. SIM images were taken of the A4-EGFP core (magenta), followed by STORM images of the L1 membrane protein (cyan). The images were aligned and overlayed. **b)** VirusMapper SPA models of A4 and L1 using the SIM/STORM data. Particles were aligned in parallel and averaged to produce the model. Scale bars 200 nm.

**Fig. S2.**
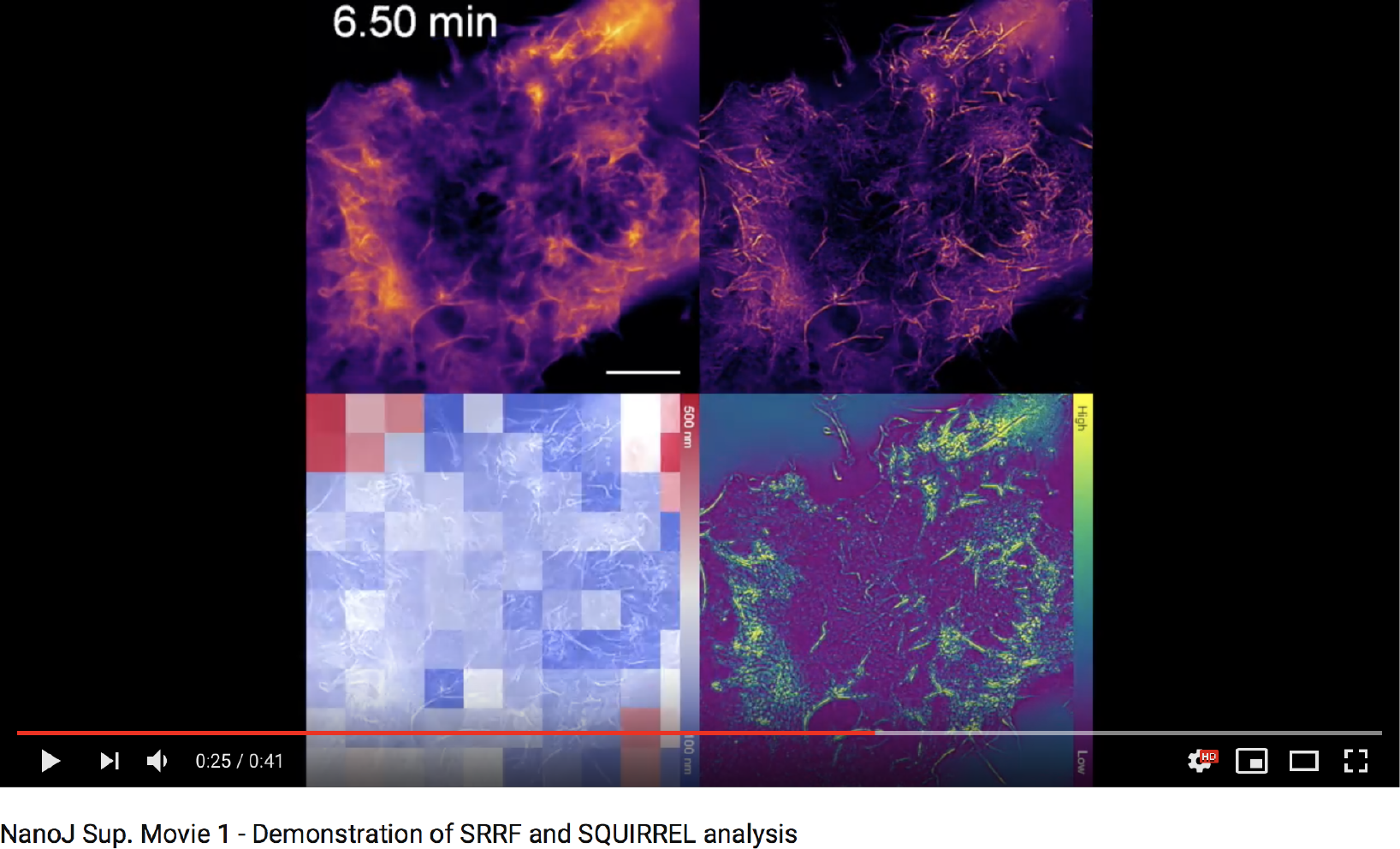
Actin dynamics visualized at super-resolution by NanoJ-SRRF. Cos7 cells expressing GFP-UtrCH imaged at 33.3 Hz, and reconstructed at 0.3Hz (100 frames) for 30 min. The image quality and resolution were assessed by NanoJ-SQUIRREL. https://www.youtube.com/watch?v=QhqKNBM4-F0feature=youtu.be Scale bars 10 μm.

